# Germline-mediated ubiquitous recombination in ScxCre male mice: implications for tendon research

**DOI:** 10.64898/2026.04.16.719028

**Authors:** Haiyin Li, Chike Cao

## Abstract

Scleraxis (Scx), a basic helix-loop-helix (bHLH) transcription factor, is a primary marker for tendon and ligament lineages. Consequently, mouse models utilizing Cre recombinase under the control of the Scx locus represents a powerful tool for control of gene expression in tendon. The constitutive ScxCre mouse line is widely used for tendon-specific genetic manipulation. In this study, we demonstrate that ScxCre exhibits undesired significant off-target activity in the male germline, leading to ubiquitous recombination of floxed alleles in all tissues of the resulting offspring. This inheritance of recombined LoxP alleles occurs independently of Cre inheritance, indicating that ScxCre-induces recombination occurs prior to meiosis in diploid germ cells. This off-target activity is not observed in female germline. These findings highlight a critical need for stringent parental sex selection when using ScxCre lines to ensure tissue-specific targeting and avoid unintentional global gene deletion or transgene activation.

## Introduction

The Cre-loxP system is one of the most powerful research tools used in mouse genetic manipulation. It enables the control of gene expression in a spatio-temporal manner and allows for cell lineage tracing by turning on the expression of a reporter transgene. This system relies on the site-specific recombinase, Cre, derived from bacteriophage P1 and two 34-bp DNA recognition sequences known as loxP sites. The Cre recombinase recognizes these sites and catalyzes the recombination of DNA sequences flanked by them (called “floxed” gene). To exploit this system, a collection of Cre mouse lines has been generated in which Cre expression is controlled by tissue-specific promoters. Crossing these Cre mice with a strain bearing a floxed target results in DNA recombination in specific cell types, yielding tissue-specific knockouts or transgene expression. The reliability of this system is critically dependent on the specificity of the chosen promoter. However, since most genes are expressed across multiple tissues, concerns persist regarding the fidelity of the Cre-loxP system and the potential for “off-target” recombination [1-5].

Scx is a basic helix-loop-helix (bHLH) transcription factor that is persistently expressed in tendons and ligaments from early developmental through adulthood [6]. Scx plays a key role in tendon progenitor cell fate, specification, differentiation and maturation [7]. Homozygous Scx^-/-^ null mice exhibit a severe loss in intermuscular tendons and those responsible for all force-transmitting, which significantly limits the use of all paws and dorsal muscles [7]. To date, Scx remains the best-characterized and most widely used marker for the tendon lineage [6, 8, 9]. Consequently, genetic manipulation of the Scx locus provides a unique opportunity to target gene expression in tendon progenitors and mature tenocytes. The constitutive ScxCre transgenic mouse line, generated by BAC transgenesis [10], is a powerful tool for modulating gene expression during tendon development and pathology.

In this study, we demonstrate that ScxCre transgene leads to the germline deletion of loxP-flanked alleles when carried in the male germline, but not the female. Consequently, offspring of ScxCre-expressing males exhibit germline-recombined alleles and subsequent gene activity ubiquitously across all tissues. Notably, even offspring that do not inherit the ScxCre transgene itself still possess the recombined alleles, indicating that Cre-medicated recombination in the male germline occurs in premeiotic diploid cells. Therefore, when ScxCre and a floxed allele are carried together in a male parent, ScxCre-mediated gene targeting loses its tissue specificity. To avoid unintended germline recombination, breeding strategies must be carefully designed; specifically, the ScxCre transgene should be introduced through the female parent to ensure the target alleles remain tissue-specific and to avoid the paternal germline leakage observed in male carriers.

## Results

### Detection of recombined loxP alleles in Cre-negative offspring of ScxCre mice

To assess the effects of tendon fibroblast-specific CaV1.2 gain-of-function in vivo, as we previously reported [11], we crossed ScxCre male mice with CaV1.2-TS^fl/fl^ mice. Resultant ScxCre;CaV1.2-TS^fl/+^ litters developed wavy tails with 100% penetrance in both male and female (Fig. 1A), a phenotype resulting from abnormal tail tendon formation induced by mutant CaV1.2^TS^ channel expression [11]. Unexpectedly, when we applied tdTomato as a lineage-tracing reporter by crossing male ScxCre;Ai9^fl/+^ with female CaV1.2-TS^fl/fl^ mice, we observed reporter activation in Cre-negative offspring: Pup1 possesses a normal, straight tail (indicating no Cre expression), however, exhibiting ubiquitous tdTomato expression, resulting in a visible red-pinkish hue in the tail, limbs and ears (Fig. 1B, Pup 1). Further, PCR genotyping of genomic DNA (gDNA) from ear biopsies confirms the absence of the ScxCre transgene in Pup 1, despite the presence of the recombined tdTomato fragment (Ai9^Δ^) was produced (Fig. 1C). These data indicate that germline recombination occurs in ScxCre mice when this breeding strategy is employed, leading to the paternal inheritance of the pre-recombined alleles.

**Figure 1.**
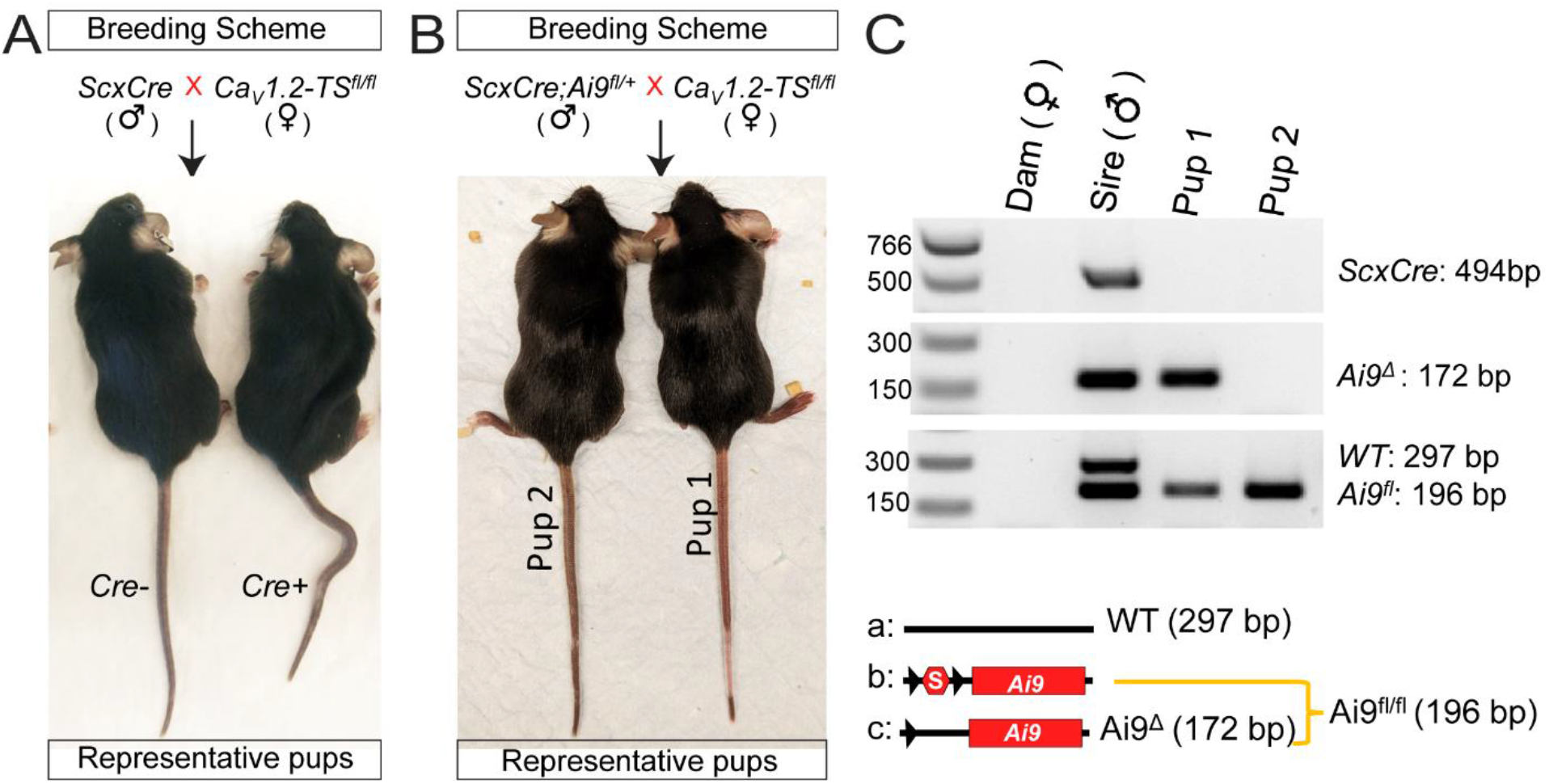
Breeding with ScxCre males causes germline recombination and unexpected recombined alleles in Cre-negative offspring. (A) Representative images of ScxCre;CaV1.2-TS^fl/+^ mice showing the wavy-tail phenotype compared with a ScxCre-negative littermate control. (B) Representative progeny generated by breeding male ScxCre;Ai9^fl/+^ mice with female CaV1.2-TS^fl/fl^ mice, demonstrating germline activation of the Ai9 reporter in offspring lacking inherited ScxCre. (C) PCR genotyping of ear-punch DNA from dam, sir and two pups using primers for ScxCre (top; 494bp), the recombined Ai9^Δ^ allele (middle; 172 bp), and the unrecombined WT (297 bp) and Ai9^fl^ (196bp) alleles (bottom)

To independently validate this observation using a different floxed allele, we crossed male ScxCre;RLacZ^fl/+^ mice with female C57BL/6J mice. The presence and absence of Cre and RlacZ transgenes in the generated offspring were determined by PCR. To detect RlacZ transgene recombination and subsequent β-galactosidase activity, whole-mount X-gal staining was performed on neonatal mice (P0-P4). While standard Mendelian inheritance predicts only four potential gentotypes (Fig. 2A, genotype 1, 2, 3 and 4), the occurrence of germline recombinaton yields additional two possible genotypic outcomes (Fig. 2A, genotype 5 and 6). As expected, no β-galactosidase activity was observed in 38% of the offspring, representing those with genotypes 1, 2, and 3 (Fig. 2B). In 19% of the progeny, Cre activity was appropriately restricted to the tendons and ligaments of the limbs, ribs, and craniofacial sutures, representing those with genotypes 4 (Fig. 2C). However, approxiamately 43% of the progeny displayed widespread X-gal staining throughout the entire body (Fig. 2D), indicating ectopic germline recombination. Notably, within this “whole-body” group, 5 out 9 pups were Cre-positive (Genotype 5), while 4 out 9 pups had not inherited the ScxCre transgene (Genotype 6). This distribution suggests that recombination occurred during gametogenesis prior to the completion of meiosis. The frequency of each observed genetoype is summarized in Table 1. Collectively, these data show that a germline-recombined gentoype occurred in 43% of all offspring sired by a ScxCre-positive male.

**Table 1:** Frequencies of offspring genotypes by parental Cre source.

| **Cre-bearing parent** | **Offspring Genotype** |  |  |  |  |  |
| --- | --- | --- | --- | --- | --- | --- |
|  | type 1 | type 2 | type 3 | type 4 | type 5 | type 6 |
| *Cre* in sire | 0/21 | 6/21 | 2/21 | 4/21 | 5/21 | 4/21 |
| *Cre* in dam | 7/22 | 4/22 | 4/22 | 7/22 | 0/22 | 0/22 |

Note: Total 21 pups from 4 independent litters for Cre in Sire, and 22 pups from 4 litters for Cre in Dam were analyzed.

**Figure 2.**
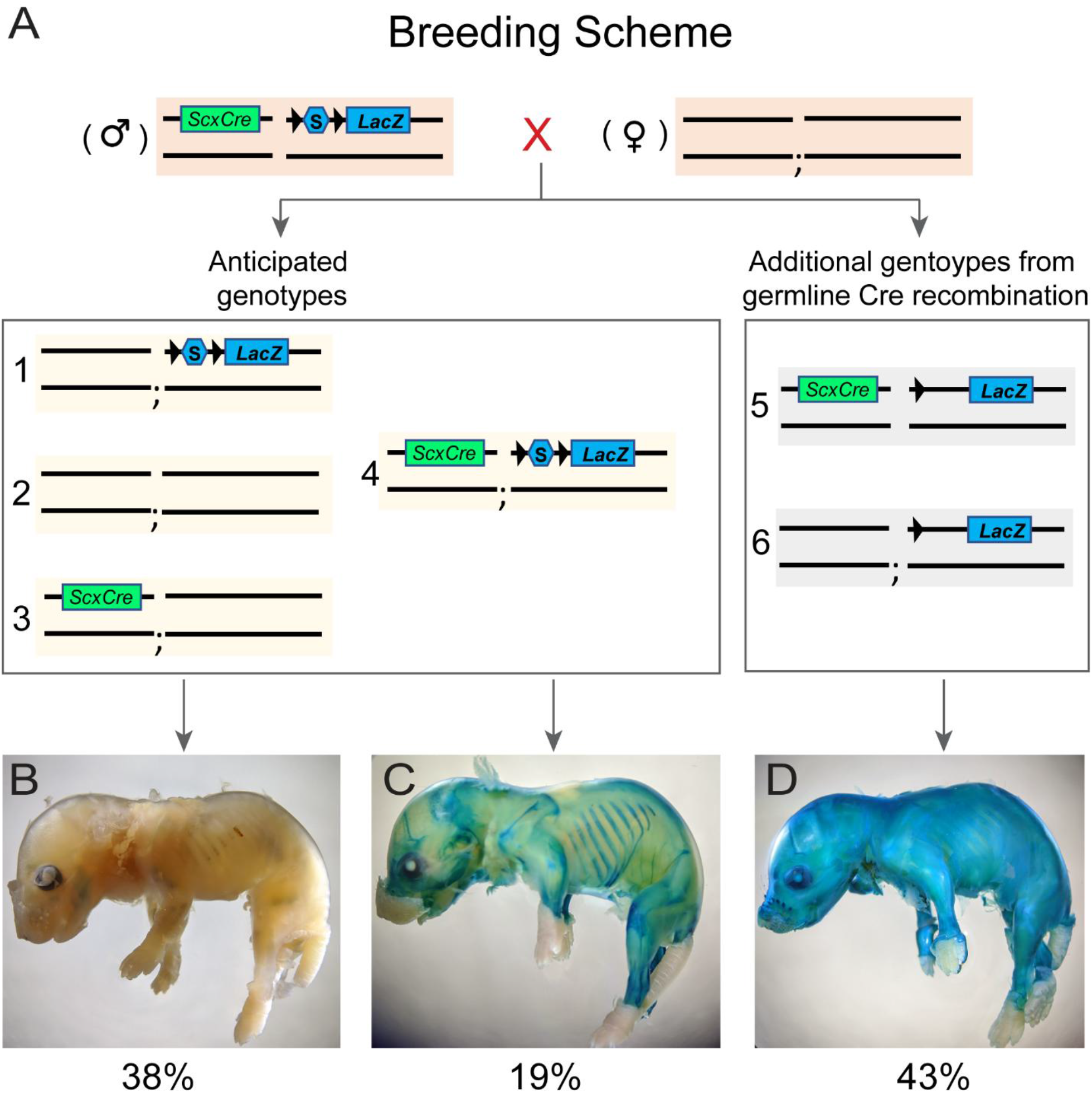
Germline recombination of the RlacZ floxed allele in male ScxCre mice. (A) Schematic showing the anticipated genotypes (type 1, 2, 3, and 4) and the additional genotypes (type 5 and 6) generated by germline Cre recombination from crosses between male ScxCre;RLacZ^fl/+^ mice and female C57B/6J mice. (B-D) Representative whole-mount X-gal staining images of offspring from this cross, showing unstained, specifically stained, and diffusedly stained patterns, respectively. Blue staining indicates β-galactosidase expression after Cre mediated recombination of the RlacZ reporter allele. The percentages under each image indicate the frequency of each staining pattern among offspring. In total, 21 pups from 4 independent litters were analyzed.

To determine whether the ScxCre transgene triggers germline recombination when transmitted maternally, we cross female ScxCre;RLacZ^fl/+^ mice with male C57BL/6J mice. In contrast to the paternal crosses, we observe no instances of widespread X-gal staining in the resulting progeny. Furthermore, PCR genotyping confirmed that β-galactosidase activity remained strictly dependent on the inheritance of the ScxCre transgene, with no evidence of pre-meiotic recombination. The observed genotype frequencies for this maternal inheritance strategy were summarized in Table 1.

## Discussion

Scx is widely regarded as a definitive marker for tendons and ligaments [6, 7, 12]. To date, the ScxCre mouse line generated by Dr. Ronen Schweitzer and colleagues remains an essential tool for manipulating gene expression during tendon/ligament development and disease pathology [10]. However, endogenous Scx expression is also present in other tissues, including testes [12-14], which raises the potential for germline recombination. In this study, we demonstrate that when ScxCre is transmitted paternally, 43% of offspring exhibit recombined RlacZ alleles in all tissues, regardless of the tissue-specific expression of Scx or the inheritance of the ScxCre transgene itself. While recombination efficiency may vary depending on the specific floxed target, this germline recombination was notably absent in the offspring of ScxCre females. This is consistent with previous findings that Scx mRNA is not detected in the ovary [13]. Based on these data, utilizing ScxCre females as the Cre-carrying parent is the necessary breeding strategy to ensure tissue-specific gene manipulation and avoid unintended global recombination.

Strong evidence indicates that endogenous Scx expression is restricted in Sertoli cells in the seminiferous tubules of the testes [13-15]. As these are supporting cells that provide nutrients to developing germ cells rather than being part of the germline itself [16, 17], this would typically argue against ScxCre activity in the male germline. However, the ScxCre mouse line was generated via BAC (bacterial artificial chromosomes) transgenesis [10]. The large BAC regulatory region used to drive tissue-specific expression may not fully recapitulate the endogenous expression of Scx, potentially leading to ectopic or “leaky” Cre activity within the germline.

Germline recombination for in many Cre lines was not recognized given the fact that screening for such events typically requires a two-generation breeding scheme. In this approach, F1 offspring (Cre^+^; reporter^fl/+^) are crossed with wild-type mice, and reporter expression is then evaluated in the F2 generation. To prevent unintended recombination and the subsequent misinterpretation of phenotypes, rigorous characterization of any Cre line is essential. First, Cre activity outside the target tissue, especially in the testes or ovaries, should be validated by crossing the Cre-driver mouse with a floxed reporter and examining the F1 generation. Second, for transgene expression studies, breeding strategies should avoid using a single parent that carries both the Cre and the floxed transgene to prevent the co-inheritance of these elements. Finally, in conditional knockout studies, additional PCR genotyping using primers that span the floxed region is mandatory to detect the presence of the recombined fragments. These steps are critical to tracking ectopic recombination and ensuring that observed phenotypes are truly tissue specific.

## Materials and Methods

### Experimental mouse models

All animal procedures were performed in accordance with the protocols approved by the Institutional Animal Care and Use Committee at the University of Rochester Medical Center. The transgenic ScxCre moue line was generously provided by Dr. Ronen Schweitzer. The ScxCre [10] and CaV1.2TS^fl/fl^ [18] mouse lines have been described previously. C57BL/6J, RlacZ^fl/fl^, Ai9^fl/fl^ strains were obtained from the Jackson Laboratory. All mice were on a C57BL/6J genetic background. To trace ScxCre and CaV1.2TS mutant channel expression, male ScxCre;Ai9^fl/+^ mice were crossed with female CaV1.2TS^fl/fl^ mice. For examining recombination in testes and ovaries, ScxCre mice were crossed with RlacZ^fl/fl^ mice to generate ScxCre;RlacZ^fl/+^ progeny, which were subsequently backcrossed with male or female C57BL/6J mice.

### X-gal staining

Whole-mount X-gal (5-bromo-4-chloro-3-indolyl-β-D-galactopyranoside) staining was performed on neonatal mice between P0 and P4. Samples were fixed in 4% paraformaldehyde on ice for 1 hour, washed three times with 1x PBS and incubated in X-gal staining solution (5mM potassium ferrocyanide, 5mM potassium ferricyanide, 1 mg/ml X-gal, 2mM MgCl2, 0.1% sodium deoxycholate, 0.2% IGEPAL CA-630) in the dark for 12 hours at 37 °C.

### Genotyping

Genomic DNA was extracted from ear punches and PCR genotyping was performed using GoTag^R^ Master Mixes (Promega, Cat#: M7122). The PCR amplification program consisted of an initial denaturation at 94ºC for 3 minutes, followed by 35 cycles of 94 ºC for 30 seconds, 61ºC for 1 minute and 72ºC for 1 minute, and a final cycle of 72 ºC for 5 minutes. DNA fragments were resolved on 2% agarose gel containing SYBR™ Safe DNA Gel Stain (ThermoFisher, Cat#:S33102) and visualized using a Bio-Rad gel documentation system. The primers used in this study were listed below:

ScxCre-F: 5’-GCAGAACCTGAAGATGTTCGC-3’

ScxCre-R: 5’-ACACCAGAGACGGAAATCCATC-3’

RosaWT-R: 5’-CGTACTGACGGTGGGAGAAT-3’

RosaWT-F: 5’-CTTTAAGCCTGCCCAGAAGA-3’

Ai9(floxed)-F: 5’-CTGTTCCTGTACGGCATGG-3’

Ai9(floxed)-R: 5’-GGCTTAAAGCAGCGTATCC-3’

Ai9^Δ^ (recombined)-F: 5’-CGTGCTGGTTATTGTGCTGT-3’

Ai9^Δ^ (recombined)-R: 5’-GCAGGTCGAGGGACCTAATA-3’

RlacZ WT-R: 5’-GGAGCGGGAGAAATGGATATG-3’

RlacZ common-F: 5’-AAAGTCGCTCTGAGTTGTTAT-3’

RlacZ(floxed)-R: 5’-GCGAAGAGTTTGTCCTCAACC-3’

## Acknowledgement

We are grateful to Dr. Ronen Schweitzer for providing the ScxCre mouse line. This work was supported by NIH/NIAMS P30AR69655 and R01AR085590.

